# Dark citations to Federal resources and their contribution to the public health literature

**DOI:** 10.1101/2023.03.26.533809

**Authors:** Jessica M. Keralis, Juan Albertorio-Díaz, Travis Hoppe

**Affiliations:** National Center for Health Statistics, Hyattsville, MD, United States

## Abstract

The term “dark citations”, which has been previously used to refer to citations of information products outside of traditional peer-reviewed journal articles, is adapted here to refer to those that are not linked to a known indexed identifier and are effectively invisible to traditional bibliometric analysis. We investigate an unexplored source of citations in the biomedical and public health literature by surveying the extent of dark citations across the U.S. government. We systematically focus on public health, quantify their occurrences across the government, and provide a comprehensive dataset for all dark citations within PubMed.

## Introduction

The U.S. federal government funds over $137 billion dollars towards basic, applied research and development [1]. The direct output of this research is often cataloged and tracked through an end product like a publication, patent, book, or clinical trial. These outputs are interconnected through their citations and provide a proxy of impact through influence. However, not all products can be readily tracked with an identifier. Federal agencies often release authoritative information through guidelines, fact sheets, manuals, web pages, and other informational products that are not always systematically indexed in a publication database but nonetheless contain vital scientific information, best practices, or policy guidance for health topics not available anywhere else [Table 1]. For agencies that frequently disseminate information using these resources, tracking their reach and usage is critical to assess impact, assign funds and develop a basic profile of their informational deliverables. Quantitative evaluation based on citations is now a widely accepted practice and is utilized by academics and other research institutions globally to assess research performance evaluation and impact assessment [2]. In addition to quantifying the influence of a given work or researcher on academic literature (e.g., by constructing a metric related directly to citations) [3], it can estimate the influence and knowledge contribution by the funding organizations. Common identifiers include the International Standard Book Number (ISBN) for books [4], the International Standard Serial Number (ISSN) for magazines and other serial publications [5], U.S. Patent and Trademark Office (USPTO) number for patents, clinical trials, digital object identifiers (DOI) [6] for peer-reviewed journal articles and other selected publications. With a representative sample of an organization’s publications’ products, bibliographic methods can quantify the impact and influence through citations and identify research communities.

**Table 1:**
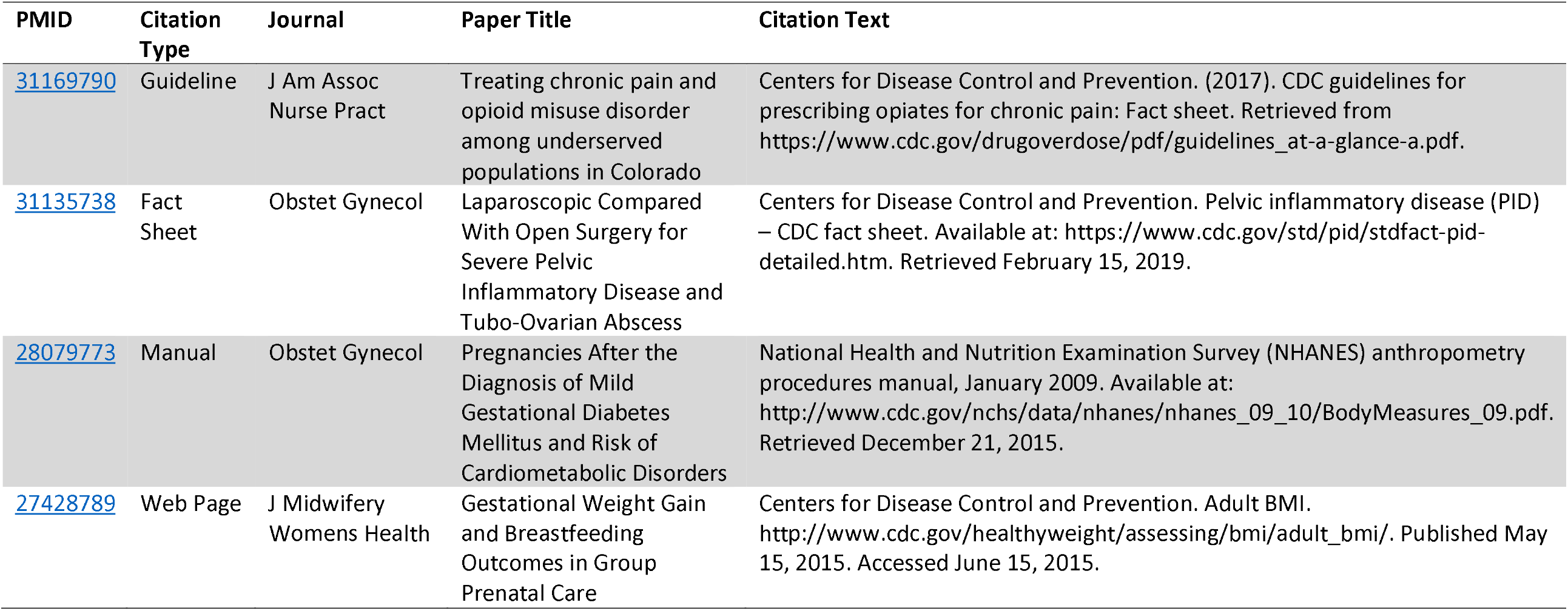
Examples of dark citations, including the source publication, journal, and citation text as it appears in the reference section. The table highlights four different types of products commonly found in dark citations: guidelines, fact sheets, manuals, and web pages.

| PMID | Citation Type | Journal | Paper Title | Citation Text |
| --- | --- | --- | --- | --- |
| 31169790 | Guideline | J Am Assoc Nurse Pract | Treating chronic pain and opioid misuse disorder among underserved populations in Colorado | Centers for Disease Control and Prevention. (2017). CDC guidelines for prescribing opiates for chronic pain: Fact sheet. Retrieved from https://www.cdc.gov/drugoverdose/pdf/guidelines_at-a-glance-a.pdf. |
| 31135738 | Fact Sheet | Obstet Gynecol | Laparoscopic Compared With Open Surgery for Severe Pelvic Inflammatory Disease and Tubo-Ovarian Abscess | Centers for Disease Control and Prevention. Pelvic inflammatory disease (PID) – CDC fact sheet. Available at: https://www.cdc.gov/std/pid/stdfact-pid-detailed.htm. Retrieved February 15, 2019. |
| 28079773 | Manual | Obstet Gynecol | Pregnancies After the Diagnosis of Mild Gestational Diabetes Mellitus and Risk of Cardiometabolic Disorders | National Health and Nutrition Examination Survey (NHANES) anthropometry procedures manual, January 2009. Available at: http://www.cdc.gov/nchs/data/nhanes/nhanes_09_10/BodyMeasures_09.pdf. Retrieved December 21, 2015. |
| 27428789 | Web Page | J Midwifery Womens Health | Gestational Weight Gain and Breastfeeding Outcomes in Group Prenatal Care | Centers for Disease Control and Prevention. Adult BMI. http://www.cdc.gov/healthyweight/assessing/bmi/adult_bmi/. Published May 15, 2015. Accessed June 15, 2015. |

As the key provider for health statistics in the United States, the National Center for Health Statistics (NCHS) has an intrinsic motivation to understand how its outputs are used. This work was motivated by the initial finding that many of the Center’s outputs, including statistics, reports, and guidelines, were cited by URL directly and were consequently difficult to track with traditional bibliometric analysis. In this work, we operationally define the term “dark citation” [3] as any citation that does not include an indexed representation. These information products, sometimes referred to as “dark data” [7], can be challenging to identify and quantify by traditional bibliometric methods. To focus on federally funded resources, we restricted our analysis to citations whose reference text included a uniform resource locator (URL) pointing to the top-level U.S. governmental domain (.gov). We evaluated all publications, federally funded or not, within PubMed that included a structured XML) reference section. The focus on PubMed allows a first look into the prevalence and nature of these dark citations in the biomedical and public health domain. Interestingly, the ability to detect dark citations within PubMed is a relatively new phenomenon. This is likely due two major contributing factors: the rise of digital-only reports published by governmental agencies and the inclusion of reference sections within PubMed that can be parsed. While nearly every publication indexed by PubMed includes a reference section, this information is not always included in a machine-readable format making systematic analysis impossible. The presence of a reference list within PubMed has changed within the last two decades: rising from about 6% in 2000 of all publications to nearly 60% in 2021 [Supplemental Figure 1].

This works aims to systematically survey the extent and quantify dark citations across the U.S. government with a focus on public health. We quantify dark citations by the most prevalent hierarchy within the U.S. government: by branch, department, agency, center, and division. We assess the health of the links within these citations to see if they still resolve to a valid webpage and discuss the implications of dark citations within bibliometric analysis. A detailed explanation of the methods and presentation of the results are available in the Supplemental Information.

## Results and Discussion

103,484 dark citations were identified among references cited by all publications for all years within the entire PubMed database; the majority were found from 2000 onward [Supplemental Figure 1]. Federal executive agency websites comprised the overwhelming majority (92%) of citations. Nearly three quarters of all federal executive dark citations came from agencies within the U.S. Department of Health and Human Services (HHS). The U.S. Centers for Disease Control and Prevention (CDC) had the largest fraction of these citations (35%), followed by the Food and Drug Administration (FDA) (26%) and the National Institutes of Health (NIH) (16%). While a large percentage of references to NIH websites were for cancer-related topics – 38% were within the cancer.gov domain – a notable number of CDC dark citations were due to references on the SARS-CoV-2 novel coronavirus and the COVID-19 pandemic. By counting subdomains like covid.cdc.gov or subdirectories like www.cdc.gov/vaccines/covid-19, the fraction of CDC’s dark citations related to COVID-19 were estimated to be at least 16%. CDC web-based resources were primarily sources of information on COVID-19 testing, case surveillance, and mortality data. In contrast, the FDA resources were focused more on authorization for the tests, treatments, and vaccines for COVID-19. Among CDC Centers, the NCHS (15.6%) and the National Center for Chronic Disease Prevention and Health Promotion (14.8%) accounted for the highest numbers of dark citations.

The importance of reference tracking in general, and dark citations specifically, varies by the entity conducting the bibliometric analysis. Peer-reviewed journals, for example, rely on the calculated impact factor [8] or other citation-based metrics [9] as the “gold standard” by which their reach and influence on research in the field are assessed and by which they are compared to other journals. Indexed identifiers (e.g., DOIs) are essential for bibliometric analyses, as nearly all peer-reviewed publications assign one to every article they publish. Organizations such as government agencies, think tanks, advocacy groups, and other non-profit entities track references to their work as a means of demonstrating their reach, influence, and value to stakeholders, particularly donors (for non-governmental organizations) or taxpayers (as represented by legislative assemblies, for government agencies). Regardless of the specific motive, tracking usage of published materials through frequency of citation is an important means of demonstrating and quantifying impact and influence for both individuals and organizations. Thus, understanding the full reach and usage of dark citations may become more necessary as such citations become more frequent.

Using bibliometrics to track references to agency websites may have greater relevance for some agencies and less for others, as some agencies’ work may not involve research publication. For example, agencies with mandates primarily related to conducting intra or extramural research such as the National Institutes of Health have a direct incentive to produce indexed products such as peer-reviewed journal articles, patents, or clinical trials [10]. In contrast, agencies such as the Centers for Disease Control and Prevention or the Food and Drug Administration have additional mandates to produce science-based practice guidelines, policy documents, recommendations, or authoritative statistics. These government-produced materials were often cited directly in the reference section and are the primary focus of this analysis.

The New England Journal of Medicine’s Ingelfinger rule - which stipulated that the journal would only consider a manuscript for publication if its substance has not been submitted or reported elsewhere [11] largely shaped traditional scientific norms around publishing in the latter half of the 20th century [12]. However, modern publishing and scientific consumption have challenged some of these norms, including the rise of preprints, social media, and web-only digital products. Rather than relying on traditional media to disseminate published findings, U.S. federal government agencies now work to make information easily accessible to the general public, 85% of whom own a smartphone [13]. Additionally, information on an official agency website is considered authoritative and accepted as a reliable source of information in scientific research, and many agencies seek to make their websites the primary source and dissemination platform for their scientists’ work.

Because the primary purpose of citations in a research manuscript is to demonstrate that the theoretical framework and methods on which the work is based is sound, and were drawn from authoritative sources, authors have little incentive to search for referenced information exclusively from indexed sources when “dark citations” such as government websites are accepted as authoritative by the scientific community. Thus, we expect these types of dark citations will only become more common unless agencies provide a means to register their outputs with an indexed representation. This could include assigning DOIs to guidelines, factsheets, or datasets and providing researchers with prominent suggested citation text near the associated resource. Finally, we found that dark citations suffer from a reproducibility crisis as well; about 10% of all dark citations were unable to be accessed anymore [Supplemental Table 2]. The ability to quantify and analyze dark citations will become increasingly important to the discipline of bibliometrics as scientific information dissemination norms continue to evolve.

## Supporting information

Supplemental Dataset: CSV table of all governmental dark citations collected

## Supplemental Information

### Methods

We downloaded the entirety of the PubMed database^1^ on June 6, 2022, in XML format, including both the baseline and update files. We merged the records keeping the most updated information for each PubMed IDentifer (PMID) resulting in 35,408,546 records. We filtered PMIDs that lacked a <referencelist> element tag, as these publications lacked reference information leaving 9,223,992 publications. For each reference, we eliminated those with element tags that included a PMID, PMCID, DOI, or PII element as these directly linked to an indexed article. The remaining references were parsed as free text and we scanned for URLs. For a reference to be considered a dark citation in this work it must include a URL with a top-level .gov domain, and not reference PubMed itself, clinicaltrials.gov, or paft.uspto.gov. For NIH, we folded domains that belonged to the agency, like cancer.gov.

For each.gov URL collected from a dark citation, we determined provenance (e.g., branch, department, State, agency, etc.) by matching the domain against the registrar of U.S. government domains^2^ provided by the Cybersecurity and Infrastructure Security Agency. To identify the status of the links, we programmatically accessed each link a total of 5 times over the course of a month. A link was considered valid if it returned a status code in the range of 2xx or 3xx at any point in the query. To reduce the burden on the target servers a HEAD request was attempted first and if the request failed it was followed by a subsequent GET request.

### Results

We examined the prevalence of dark citations across the biomedical literature at multiple levels of the U.S. Federal government by branch, department, agency, center, and division.^3^ We focused on the most frequent area at each level of the hierarchy: the Executive Branch (92%), the Department of Health and Human Services (HHS) (74%)^4^, the Centers for Disease Control (CDC) (35%), and the National Center for Health Statistics (NCHS) (15%). Respectively, each level of the hierarchy is represented in the following tables [Supplemental Table 3] [Supplemental Table 4], [Supplemental Table 5], [Supplemental Table 6], [Supplemental Table 7]. There were a small number of dark citations to domains operated by federally recognized tribal nations, including navajo-nsn.gov (Navajo Nation), cdatribe-nsn.gov (Coeur d’Alene Tribe), hopi-nsn.gov (Hopi Tribe), and menominee-nsn.gov (the Menominee Indian Tribe of Wisconsin). However, it is important to note that many tribes have websites within other domains, including the commercial.com domain (e.g., Eastern Band of Cherokee Indians at ebci.com or the Comanche Nation at comanchenation.com) or the non-profit.org domain (e.g., the Apache Tribe of Oklahoma at apachetribe.org).103,484 dark citations were identified in the biomedical literature whose top-level domain pointed to a .gov. Approximately 94% of these dark citations originated from the Federal level, primarily in the Executive branch 92%, 2% in the Legislative branch, and only 49 total (< 0.1%) from the Judicial branch. Four percent were found at the state level. The remaining dark citations were found in municipal, country, tribal top-level domains. A handful of dark citations were found in true multi-level domains (e.g., the Appalachian Regional Commission, www.arc.gov) or National Labs (e.g., the Ames National Laboratory at Iowa State University, www.ames.gov).

As a result of focusing on biomedical literature, it was unsurprising to find the dark citations concentrated around agencies devoted to providing guidelines and public health advice to public. HHS accounted for 74.3% of the total dark citations from the Executive branch while the Department of Commerce (5.5%), Environmental Protection Agency (3.7%), United States Department of Agriculture (2.7%), the Department of Labor (25) and the Veterans Affairs (1.8%). The remaining departments accounted for less than 2% of the total.

Focused on HHS, the Centers for Disease Control and Prevention (35%), the Food and Drug Administration (FDA) (25%), National Institutes of Health (NIH) (16%) and Center for Medicare & Medicaid Services (CMS) (6.3%) accounted for majority of the totals.

We note that dark citations arise primarily by work produced via intramural funding (as opposed to the extramural funding through grants given by the agency).

Drilling into the specifics of the dark citations from the CDC, we allocated the totals by Center, Institute, Office or data collection. Notably, a large body of public health information came recently from COIVD-19 related tables, guidelines, and reports that were not linked to a specific center (16.5%). The Centers with the largest fraction included the National Center for Health Statistics (NCHS) (15.6%), the National Center for Chronic Disease Prevention and Health Promotion (14.8%), and the National Center for HIV, STD, and TB Prevention (10.8%). The breakdown for the remaining all accounted for < 10% individually.

Finally, within NCHS, the largest divisions included the Health and Nutrition Examination Survey (29.5%) and Vital Statistics (16%), with the remaining divisions at < 10%. For this section, we noted the distinction between formal reports, such as National Health Statistics Reports, National Vital Statistics Reports, and Health E Stats, and non-report web-based resources, such as web pages and manuals. Many NCHS reports only exist as digital links, leaving research unable to cite anything except for the URL.

In general, we found that most of these links to still be valid returning an HTTP status code of in the range of 2xx (successful) or 3xx (redirection). Out of the remaining links 5.5% returned status code 404 (not found), or another other client / server errors 4xx (client) / 5xx (server) error collectively at 4.5% [Supplemental Table 2].

### Supplemental Figures and Tables

**Supplemental Figure 1:**
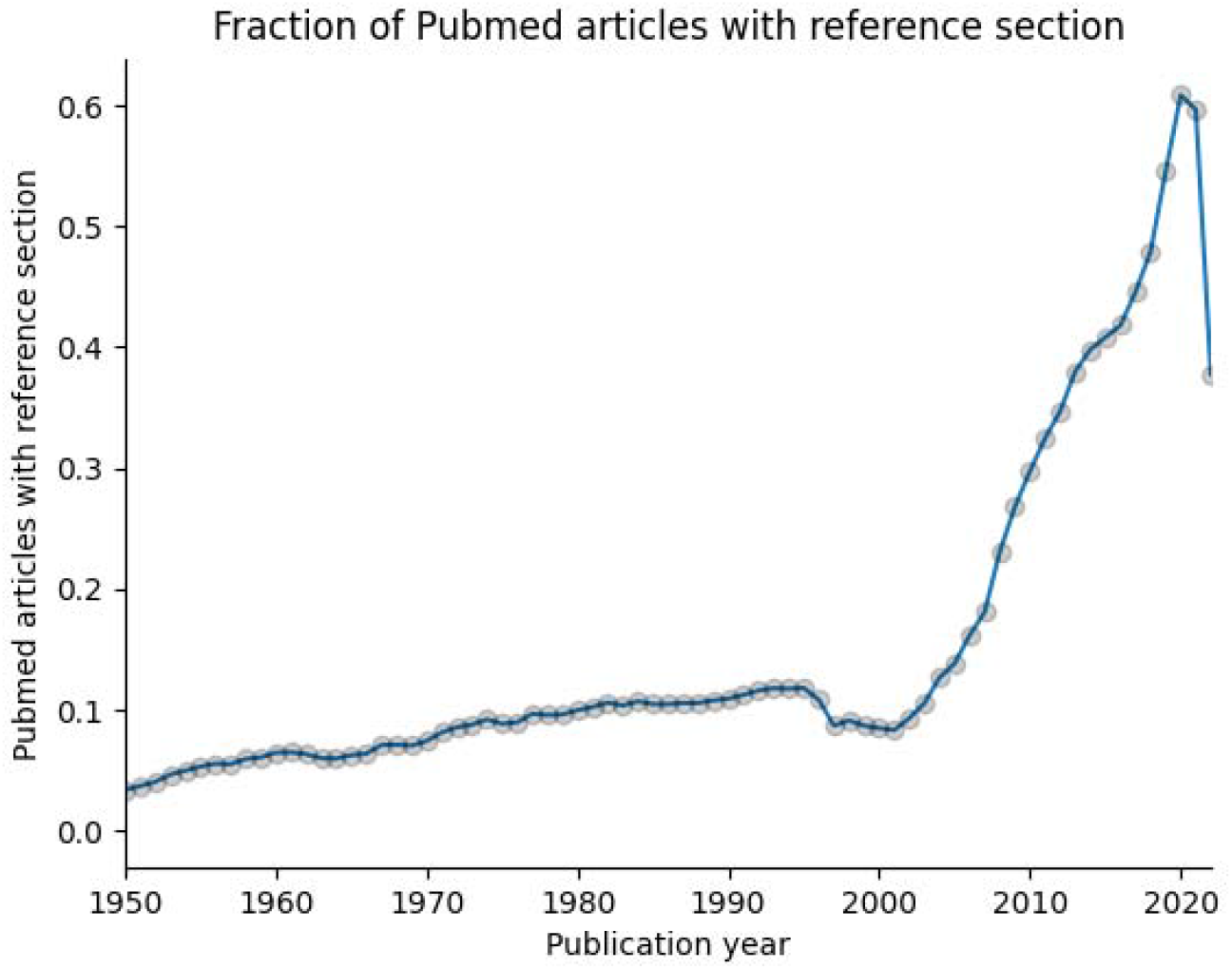
Fraction of PubMed articles with a parsed machine-readable reference section. Prior to the year 2000, less than ten percent of all publications can be parsed and analyzed. By 2018, about 60% of all articles could be analyzed. The dip in reference sections of publications in the year of this analysis (2022) is likely caused by incomplete information from publishers on new publications and may resolve in the subsequent year as records are updated.

**Supplemental Table 2:** Number and percent on the quality of dark citations. Each URL in the citation was attempted at least five times. A URL was considered valid if it returned a status code that indicated success or redirection (explicitly those in range of 200-399). This represents an upper bound on the percentage of valid dark citations, as the quality of the link or redirection was not evaluated.

| Status code returned | # of dark citations | Percent |
| --- | --- | --- |
| Valid (2xx, 3xx) | 96,690 | 90.1% |
| 404 errors | 5,862 | 5.5% |
| Client or server errors | 4,789 | 4.5% |

**Supplemental Table 3:** Number and percent of dark citations from PubMed identified across the US Federal, State, local, and tribal governments that include a .gov URL.

| **U.S. Government Level** | **# of dark citations** | **Percent** |
| --- | --- | --- |
| Federal, Executive | 96,167 | 92.1% |
| State | 4,604 | 4.4% |
| Federal, Legislative | 1,980 | 1.9% |
| Other: National Labs | 661 | 0.6% |
| Multi-level | 72 | 0.1% |
| Municipal | 662 | 0.6% |
| County | 181 | 0.2% |
| Federal, Judicial | 49 | 0.0% |
| Tribal | 9 | 0.0% |

**Supplemental Table 4:** Number and percent of dark citations from PubMed within the Federal executive branch, by Department.

| **Department** | **Acronym or shorthand** | **# of dark citations** | **Percent** |
| --- | --- | --- | --- |
| Department of Health and Human Services | HHS | 71,664 | 74.3% |
| Department of Commerce | Commerce | 5,329 | 5.5% |
| Environmental Protection Agency | EPA | 3,602 | 3.7% |
| United States Department of Agriculture | USDA | 2,618 | 2.7% |
| Department of Labor | Labor | 1,932 | 2.0% |
| Veteran Affairs | VA | 1,694 | 1.8% |
| Department of Justice | DOJ | 1,340 | 1.4% |
| Other | Other | 1,258 | 1.3% |
| Department of Education | Education | 1,209 | 1.3% |
| Department of Interior | Interior | 1,091 | 1.1% |
| Department of Energy | Energy | 902 | 0.9% |
| National Archives and Records Administration | NARA | 805 | 0.8% |
| National Aeronautics and Space Administration | NASA | 602 | 0.6% |
| Department of Homeland Security | Homeland Security | 369 | 0.4% |
| United States Agency for International Development | State/USAID | 333 | 0.3% |
| Department of Transportation | Transportation | 317 | 0.3% |
| General Service Administration | GSA | 291 | 0.3% |
| Educational Opportunity Program | EOP | 287 | 0.3% |
| Interagency |  | 275 | 0.3% |
| Department of Housing and Urban Development | HUD | 99 | 0.1% |
| Department of Treasury | Treasury | 88 | 0.1% |
| Department of Defense | DOD | 62 | 0.1% |

**Supplemental Table 5:** Number and percent of dark citations from PubMed within the US Department of Health and Human Services (HHS), by agency.

| HHS Agency | Acronym | # of dark citations | Percent |
| --- | --- | --- | --- |
| Centers for Disease Control and Prevention | CDC | 25,314 | 35.32% |
| Food and Drug Administration | FDA | 18,461 | 25.76% |
| National Institutes of Health | NIH | 11,404 | 15.91% |
| Center for Medicare & Medicaid Services | CMS | 4,515 | 6.30% |
| Health & Human Services Office of the Secretary | OS | 4,082 | 5.70% |
| Agency for Healthcare Research and Quality | AHRQ | 3,723 | 5.20% |
| Health Resources and Services Administration | HRSA | 1,944 | 2.71% |
| Substance Abuse and Mental Health Services Administration | SAMHSA | 1,918 | 2.68% |
| Administration of Community Living | ACL | 99 | 0.14% |
| Indian Health Service | IHS | 73 | 0.10% |
| Administration for Strategic Preparedness & Response | ASPR | 71 | 0.10% |
| Administration for Children and Family | ACF | 60 | 0.08% |

**Supplemental Table 6:** Number and percent of dark citations from PubMed within the US Centers for Disease Control and Prevention (CDC), by center, office, or institute.

| CDC Center, Institute, Office, system or publication set | Acronym | # of dark citations | Percent |
| --- | --- | --- | --- |
| Coronavirus Disease | COVID | 4,147 | 16.58% |
| National Center for Health Statistics | NCHS | 3,907 | 15.62% |
| National Center for Chronic Disease Prevention and Health Promotion | NCCDPHP | 3,698 | 14.78% |
| National Center for HIV, STD, and TB Prevention | NCHHSTP | 2,700 | 10.79% |
| National Center for Emerging and Zoonotic Infectious Diseases | NCEZID | 2,206 | 8.82% |
| National Center for Injury Prevention and Control | NCIPC | 1,843 | 7.37% |
| National Center for Immunization and Respiratory Diseases | NCIRD | 1,338 | 5.35% |
| Morbidity and Mortality Weekly Report | MMWR | 886 | 3.54% |
| Agency for Toxic Substances and Disease Registry | ATSDR | 680 | 2.72% |
| National Institute for Occupational Safety and Health | NIOSH | 639 | 2.55% |
| National Center on Birth Defects and Developmental Disabilities | NCBDDD | 451 | 1.80% |
| Office of the Director | OD | 438 | 1.75% |
| Wide-ranging ONline Data for Epidemiologic Research | WONDER | 351 | 1.40% |
| National Center for Environmental Health | NCEH | 281 | 1.12% |
| CDC Library | Library | 270 | 1.08% |
| Center for Preparedness and Response | CPR | 259 | 1.04% |
| Center for Surveillance, Epidemiology, and Laboratory Services | CSELS | 243 | 0.97% |
| Center for Global Health | CGH | 242 | 0.97% |
| Office of Minority Health and Health Equity | OMHHE | 101 | 0.40% |
| File Transfer Protocol | FTP | 93 | 0.37% |
| Office of the Secretary | OS | 75 | 0.30% |
| Center for State, Tribal, Local, and Territorial Support | CSTLTS | 69 | 0.28% |
| Socrata (data.cdc.gov backend) | Socrata | 46 | 0.18% |
| Severe acute respiratory syndrome | SARS | 20 | 0.08% |
| Deputy Director Public Health Science and Surveillance | DDPHSS | 17 | 0.07% |
| Office of Laboratory Science and Safety | OLSS | 11 | 0.04% |
| Occupational and Public Health Specialty Section | OPHSS | 8 | 0.03% |

**Supplemental Table 7:** Number and percent of dark citations from PubMed within National Center for Health Statistics (NCHS), by division or office split into reports and non-reports like webpages and other resources.

| Division | Acronym | # of dark citations | Percent |
| --- | --- | --- | --- |
| **Web pages and non-report resources** |  |  |  |
| Division of Health and Nutrition Examination Survey | DHANES | 1,150 | 29.49% |
| Division of Vital Statistics | DVS | 621 | 15.92% |
| Division of Health Care Survey | DHCS | 296 | 7.59% |
| FastStats (cross-divisional) |  | 262 | 6.72% |
| Division of Health Interview Survey | DHIS | 246 | 6.31% |
| Division of Analysis and Epidemiology | DAE | 204 | 5.23% |
| Research Data Center | RDC | 12 | 0.31% |
| Office of the Director | OD | 11 | 0.28% |
| Office of Information Technology | OIT | 5 | 0.13% |
| Division of Research Methodology | DRM | 3 | 0.08% |
| **Reports: NHSRs, NVSRs, SRs, data briefs** |  |  |  |
| Division of Vital Statistics | DVS | 436 | 11.18% |
| Division of Health and Nutrition Examination Survey | DHANES | 248 | 6.36% |
| Division of Analysis and Epidemiology | DAE | 92 | 2.36% |
| Division of Health Care Survey | DHCS | 60 | 1.54% |
| Division of Health Interview Survey | DHIS | 59 | 1.51% |
| Division of Research Methodology | DRM | 1 | 0.03% |
| Health E-stats |  | 79 | 2.03% |
| Press releases |  | 58 | 1.49% |
| Other/Unknown |  | 57 | 1.46% |

## Footnotes

1 *https://ftp.ncbi.nlm.nih.gov/*

2 *https://home.dotgov.gov/data/*

3 *An overview of how the U.S. federal government is organized is available at https://www.usa.gov/branches-of-government.*

4 *A listing of agencies within the U.S. Department of Health and Human Services, which is a cabinet-level federalexecutive agency, can be found at https://www.usa.gov/federal-agencies/u-s-department-of-health-and-human-services.*

